# Honeybees show an increased preference for dietary alcohol when parasitized

**DOI:** 10.1101/2025.05.30.656786

**Authors:** Monika Ostap-Chec, Weronika Antoł, Daniel Bajorek, Daniel Stec, Krzysztof Miler

## Abstract

Parasitic infections often alter host behavior, including foraging and the consumption of bioactive substances. In honeybees (*Apis mellifera*), infection with the common gut parasite *Nosema ceranae* causes metabolic disruption and increased mortality. Ethanol is a naturally occurring bioactive compound found in nectar, and honeybees exhibit high tolerance and resilience to chronic exposure. However, whether bees actively use ethanol during infection remains unclear. Here, we investigated whether *N. ceranae*-infected honeybees alter their ethanol consumption. In a feeding experiment, infected and uninfected bees were given a choice between plain sucrose solution and ethanol-spiked food (0.5% or 1% ethanol). We measured food consumption, survival, and spore load. Although overall food intake did not differ between groups, infected bees consumed a significantly higher proportion of ethanol-spiked food. Survival analysis showed that a diet containing 1% ethanol caused higher mortality than a diet containing 0.5% ethanol; however, among bees on a 1% ethanol diet, this negative effect was less pronounced in infected individuals than in controls. Spore load did not differ between treatments. These results suggest that *N. ceranae* infection induces a shift in feeding behavior towards increased ethanol intake, which may benefit infected bees by reducing mortality. This may reflect a self-medication response, although alternative explanations - such as parasite-induced manipulation or ethanol-induced changes in host physiology and immunity - remain possible. Further research into ethanol’s effects on *Nosema* spores is needed. Nonetheless, our findings provide insights into honeybee interactions with bioactive compounds and suggest that ethanol may be a behaviorally relevant dietary substance.

## Introduction

Parasites are among the most ubiquitous organisms on Earth, imposing significant fitness costs on their hosts (Dobson et al. 2008). Throughout coevolution, hosts have developed various defensive strategies to combat parasitic infections. While responses to ectoparasites involve a range of behavioral adaptations, combating endoparasites, particularly gut parasites, almost always requires some form of dietary changes. These strategies vary from alterations in food intake, diet preferences, or foraging behavior (Bernardo and Singer 2017; Schmid-Hempel 2021; Cotter and Al Shareefi 2022) to more advanced mechanisms, such as self-medication, in which infected individuals actively seek out substances that, although harmful to healthy organisms, possess therapeutic properties (de Roode et al. 2013; Abbott 2014). Such behavioral responses are not always adaptive; they may also result from parasite manipulation or be unintended by-products of the infection (Vale et al. 2018). Dietary changes influenced by parasitic infection have been documented across various taxa, including insects. For instance, infected caterpillars opt for diets higher in protein and lower in carbohydrates, which increases their chances of survival (Povey et al. 2014). Honeybees actively select foods with antibiotic properties when infected (Gherman et al. 2014; Pusceddu et al. 2019). Infected ant colonies have also been observed to choose a diet that, while detrimental to long-term survival, provides short-term benefits in fighting infection (Csata et al. 2024).

Honeybees are one of the most important pollinators for both natural ecosystems and agriculture (Klein et al. 2018; Papa et al. 2022) and face various parasites. *Nosema ceranae* and *N. apis* are two extensively studied microsporidia due to their commonness and significant impact on honeybee health (Moritz et al. 2010; Hristov et al. 2020; Ostap-Chec et al. 2024a). In the external environment, they exist as spores, which must be eaten by a bee, either through food or water, for the infection to initiate (Fries 2010; Smith 2012). Spores can also be transferred sexually between drones and queens (Roberts et al. 2015). When the spores reach the midgut, they invade the epithelial cells and rapidly multiply, consuming the cell’s contents through phagocytosis and depleting the host’s resources (Gisder et al. 2011). The spores released upon cell destruction can infect other cells or be expelled, contaminating floral resources and the nesting environment. Over time, this cell destruction leads to gut lesions, impaired nutrient absorption, and other detrimental effects (Goblirsch 2018; Paris et al. 2018).

Of these two *Nosema* species, *N. ceranae* has garnered more attention due to its wider host range, ability to infect year-round with minimal seasonal variation, and higher biotic potential across different temperatures (Martín-Hernández et al. 2009; Higes et al. 2010). Unlike *N. apis*, *N. ceranae* does not cause conspicuous disease symptoms such as dysentery. However, infected bees still suffer from digestive disorders, lethargy, and a shortened lifespan (Higes et al. 2008; Higes et al. 2009; Higes et al. 2010; Botías et al. 2013; Koch et al. 2017; Ostap-Chec et al. 2024a). *N. ceranae* infection disrupts energy metabolism in honeybees, as evidenced by the dysregulated expression of metabolic genes (Vidau et al. 2014; Kurze et al. 2016). This metabolic stress is accompanied by significant immunosuppression (Antúnez et al. 2009; Chaimanee et al. 2012; Holt et al. 2013; Huang et al. 2016; Li et al. 2018; Lourenço et al. 2021). Several studies have reported a decline in hemolymph trehalose – a key energy-storage sugar – in infected honeybees (Blatt and Roces 2001; Thompson 2003; Mayack and Naug 2010; but see Ostap-Chec et al. 2025). Infected bees demonstrate increased respirometric activity and lipid loss (Li et al. 2018), ultimately leading to fat body depletion (Gilbert et al. 2024). These metabolic disruptions are further facilitated by the fact that *Nosema* spores lack mitochondria and rely entirely on the host for essential energy resources (Tsaousis et al. 2008). Such energetic strains manifest not only in physiological impairments, including disrupted digestion, thermoregulation, and increased susceptibility to starvation (Mayack and Naug 2009; Martín-Hernández et al. 2011; Vidau et al. 2014), but also in behavioral changes, such as prolonged foraging times and reduced flight frequency (Dussaubat, Sagastume, et al. 2013; Alaux et al. 2014; Naug 2014; Wolf et al. 2014). These effects can impact the entire colony, ultimately reducing its success (Emsen et al. 2020).

How honeybees combat *Nosema* infections remains insufficiently studied. Behavioral avoidance appears particularly challenging, as the disease primarily spreads through fecal-oral transmission and contaminated floral resources (Fries 2010; Smith 2012). Studies show that infected bees exhibit precocious and increased flight activity, which may help reduce pathogen transmission while keeping healthy bees engaged in safer hive tasks and reducing pathogen transmission inside the hive (Dussaubat, Maisonnasse, et al. 2013). Moreover, *Nosema*-infected bees prefer honey with higher antibiotic activity, which lowers microsporidian loads (Gherman et al. 2014). Surprisingly, despite the significant energetic stress caused by *Nosema* infection, bees do not compensate for these losses by increasing sucrose consumption (Ostap-Chec et al. 2025). Propolis, a natural hive component, is known to reduce mortality, infection rates, and pathogen infectivity in *Nosema*-infected bees (Naree, Benbow, et al. 2021; Naree, Ellis, et al. 2021). However, there is no evidence that honeybees actively employ it to fight *Nosema* (Mura et al. 2020).

Ethanol is another naturally occurring substance that may serve as a potential medicinal compound. Ethanol is widespread and found in various ecological contexts. It is produced through the fermentation of sugars, mainly by yeast (such as *Saccharomyces cerevisiae*), a process that has likely been occurring for around 100 million years (Thomson et al. 2005). With over 325,000 species of flowering plants globally, providing sugar-rich substrates (Govaerts et al. 2021), and the fact that yeasts are frequently present in nectar (Lievens et al. 2015), ethanol appears to be naturally present in nearly every ecosystem. While it was found that in tropical regions ethanol concentration in palm flower nectar can reach 6.9% (Goodrich et al. 2006; Hockings et al. 2015), in temperate conditions, ethanol concentrations are unlikely to reach more than 1% (Ehlers and Olesen 1997; Jakubska 2005; Goodrich et al. 2006; Wiens et al. 2008; Rering et al. 2018). There is growing evidence that ethanol is ecologically relevant and has shaped the evolution of many species, influencing nutritional, medicinal, and behavioral traits not only in frugivorous vertebrates (Bowland et al. 2025) but also in nectarivorous invertebrates such as bees and wasps (Miler 2025). In honeybees, ethanol plays a crucial role in the biosynthesis of ethyl oleate, a pheromone that regulates colony demography by preventing the transition of workers into foragers (Leoncini et al. 2004; Castillo, Chen, et al. 2012; Castillo, Maisonnasse, et al. 2012). Furthermore, foragers, which leave the hive to search for food, produce alcohol dehydrogenase enzymes that likely break down ethanol (Miler et al. 2021).

Honeybees willingly consume ethanol dissolved in sucrose under both laboratory and field conditions, even at high concentrations reaching up to 20% (Abramson et al. 2000; Abramson, Sheridan, et al. 2004; Maze et al. 2006; Sokolowski et al. 2012; Mustard et al. 2019). Moreover, foragers prefer feeding solutions containing up to about 2.5% ethanol over pure sucrose solutions (Mustard et al. 2019). However, the effects of ethanol on honeybees remain insufficiently explored. Early studies primarily focused on single ethanol exposure, often using high, ecologically irrelevant doses, revealing that such acute consumption impairs honeybee physiology and behavior. Ethanol disrupts social communication among workers by impairing antennation and trophallaxis (Mixson et al. 2010; Wright et al. 2012), alters dance communication (Bozic et al. 2006), and increases aggression (Abramson, Place, et al. 2004; Ammons and Hunt 2008). Additionally, it negatively affects locomotion, foraging efficiency, and learning abilities (Abramson et al. 2005; Maze et al. 2006; Mustard et al. 2008; Giannoni-Guzmán et al. 2014; Black et al. 2021; Ahmed et al. 2022), with the severity of these impairments increasing in a dose-dependent manner (Bozic et al. 2006; Maze et al. 2006; Wright et al. 2012). In natural conditions, it is more realistic that bees encounter ethanol in relatively low concentrations but repeatedly. Studies that simulate such conditions show that bees exhibit tolerance, showing reduced motor impairment compared to naïve individuals (Miler et al. 2018; but see Stephenson et al., 2021). Chronic dietary ethanol exposure, whether through voluntary consumption of 0.5% ethanol or forced intake of 1%, has no significant effect on survival, flight endurance, body mass, or lipid content, except for an increased trehalose level in the hemolymph (Ostap-Chec et al. 2024b). These findings suggest that honeybees are highly resilient to chronic intake of ethanol. However, research on older bees indicates that both occasional and continuous ethanol exposure significantly reduce survival, suggesting that the toxic effects of even low concentrations of ethanol may be more pronounced in aged individuals (Ostap-Chec et al. 2024c). Interestingly, honeybees subjected to prolonged ethanol consumption exhibit withdrawal symptoms upon discontinuation of access to ethanol-spiked food (Ostap-Chec et al. 2021), a hallmark characteristic of alcohol dependence.

Ethanol is a substance that, while detrimental in high doses, may have protective or otherwise beneficial effects at low concentrations. There are examples in the animal kingdom where ethanol is used for self-medication or to enhance reproductive success. *Drosophila melanogaster* preferentially oviposits on ethanol-containing substrates, which enhance offspring fitness (Bokor and Pecsenye 2000; Azanchi et al. 2013). Additionally, *D. melanogaster* larvae increase ethanol intake when parasitized by endoparasitoid wasps, suggesting a form of self-medication (Milan et al. 2012). A comparable ethanol-based mechanism has been observed in ambrosia beetles (*Xylosandrus germanus*), which rely on ethanol-rich environments to cultivate fungal gardens essential for their reproduction (Ranger et al. 2018). As ethanol is also a caloric substance, it may serve as an alternative energy source. Butterflies appear to be physiologically adapted to ethanol, as low concentrations do not impair their fitness traits (Miller 1997). They have evolved tolerance mechanisms allowing them to derive comparable energetic value from ethanol as from sugar, which may support fecundity in conditions of limited sugar availability (Beaulieu et al. 2017). A similar strategy is observed in the desert-adapted *D. mojavensis*, where exposure to ethanol vapor increases longevity and fecundity under carbohydrate-poor conditions, suggesting that ethanol can function as an alternative energy source (Etges and Klassen 1989). This effect is genotype- and environment-dependent and likely reflects local adaptations (Starmer et al. 1977). Nevertheless, despite the frequent environmental exposure of honeybees to ethanol, its potential adaptive significance –whether medicinal, protective, or metabolic – remains insufficiently studied.

Here, we tested the hypothesis that honeybees alter their ethanol intake when infected with *Nosema ceranae*. This addresses key questions in behavioral ecology: whether hosts adaptively shift their behavior under parasitic stress and what types of responses such shifts represent. We conducted a feeding experiment that mimicked a naturalistic choice scenario between ecologically relevant, low ethanol concentrations. Healthy and *Nosema*-infected bees were offered a choice between plain sucrose solution and either 0.5% or 1% ethanol-spiked sucrose, and we measured food consumption, survival, and spore load. Specifically, we asked whether (1) infected bees increase their intake of ethanol-spiked food compared to uninfected bees, (2) ethanol consumption alters mortality depending on infection status, and (3) parasite loads differ between ethanol concentrations. Based on prior evidence of high ethanol resilience in honeybees (Ostap-Chec et al. 2024b) and infection-induced changes in dietary preferences in other systems (Milan et al. 2012; Gherman et al. 2014; Csata et al. 2024), we predicted an increase in ethanol consumption among infected bees, and a possible reduction in infection-related costs (e.g., mortality and/or spore load) due to this dietary shift.

## Methods

### Experimental procedure

The experiment was conducted using queen-right colonies of *Apis mellifera carnica* with naturally inseminated queens. The colonies were in good overall condition and had been treated with oxalic acid against *V. destructor* in early spring. Additionally, all colonies were screened for background *Nosema* infection (see below).

The study was performed in four replicates, with a two-day interval between each replicate. For each replicate, newly emerged bees were obtained from two unrelated colonies. To achieve this, a brood frame with capped cells, free of adult bees, was collected from each colony and placed overnight in an incubator (KB53, Binder, Germany) set at 32 °C. The following morning, all newly emerged bees were marked with a colored dot on the thorax using a non-toxic paint marker and introduced into an unrelated hive for 7 days. This step allowed the bees to develop in a natural hive environment, supporting physiological development, enhancing immune function, and increasing survival rates in subsequent laboratory conditions (Ostap-Chec et al. 2021, 2024c).

At seven days old, the marked bees were recollected from the hive using entomological forceps, transferred to wooden cages, and transported to the laboratory for individual feeding. To enhance feeding motivation, the bees were food-deprived for approximately one hour before treatment. Each bee was then placed in a plastic Petri dish with a lid (diameter 5.5 cm) and assigned to one of two groups: infected or control. Bees in the infected group received a 10 µL droplet of 1 M sucrose solution containing 100,000

*N. ceranae* spores (for details on solution preparation see below), while control bees received a 10 µL droplet of 1 M sucrose solution without spores. The bees were observed for up to three hours, and only those that fully consumed their respective solutions were included in the experiment.

For each replicate, 20 cages were established, 10 for infected bees and 10 for control bees, each containing 40 individuals. Over the following two days, all bees received a 40% sucrose solution and water, provided *ad libitum* via gravity feeders. The cages were maintained in an incubator (KB400, Binder, Germany) at 32 °C. After this acclimation period, the initial number of live bees in each cage was recorded. This period allowed the bees to recover from handling stress, stabilize their feeding behavior, and minimize bias from initial mortality.

After acclimation, bees within each group (infected vs. control) entered the experimental phase and were assigned to one of two dietary treatments. In each treatment, bees were provided with two feeders: one containing pure 1 M sucrose solution and the other containing sucrose solution with either 0.5% ethanol [diet: 0.5% EtOH] or 1% ethanol [diet: 1% EtOH]. Each dietary treatment was replicated in five cages per group. The ethanol-supplemented solutions were prepared to be iso-caloric with the 1 M sucrose solution by reducing the sucrose content by 1.38 g for each mL of ethanol (assuming 5.5 kcal per 1 mL of ethanol) to prevent confounding effects related to ethanol’s caloric value when digested (Miler et al. 2021).

The bees remained on their assigned diets for 15 days. This period reflects the upper-level longevity of forager honeybees under natural conditions and thus corresponds to the typical duration of potential ethanol exposure in the wild (Dukas 2008). Every three days, both feeders in each cage were weighed before and after refilling to assess daily food consumption. As a result, food consumption was measured at five timepoints (on days 3, 6, 9, 12, and 15 of the experimental phase, corresponding to days 12, 15, 18, 21, and 24 of the bees’ life). Mortality was recorded daily, and dead individuals were removed. In total, 3200 bees were used in the experiment (4 replicates × 2 groups × 2 diets × 5 cages × 40 individuals).

On the final day of the experimental phase, two bees from each cage were collected and frozen for subsequent molecular analysis. Quantitative PCR (qPCR) was performed to confirm infection status (infected vs. control).

### Preparation of spores for infection

We sourced the spores from our stock population of infected honeybees, which were maintained in controlled incubator conditions to sustain the infection for spore harvesting. The identity of *N. ceranae* spores in the stock population was confirmed as described by Berbeć et al. (2022). The spore suspension used for experimental inoculation was freshly prepared on the same day the bees were fed. To prepare the suspension, we homogenized the digestive tracts of several infected individuals using a micropestle in distilled water. The mixture was centrifuged (Frontier 5306, Ohaus, Switzerland) at 6,000 G for 5 minutes, repeating this process three times. After each centrifugation, the supernatant was replaced with fresh distilled water.

The final supernatant was replaced with a 1M sucrose solution, and the concentration of spores was determined using a Bürker hemocytometer under a Leica DMLB light microscope equipped with phase contrast (PCM) and a digital camera. The final infection solution was adjusted to achieve a concentration of 100,000 spores per 10 µl by diluting with 1M sucrose solution.

### Verification of *Nosema* infection status

To confirm that bees exposed to *N. ceranae* spores were indeed infected and that bees in the control group remained uninfected, we assessed spore levels using qPCR quantification. We sampled 10 infected and 10 control bees per colony and dietary treatment (160 individuals in total).

Additionally, to ensure that the colonies used for the experiment were initially free from *N. ceranae* infection, we tested their infection status. For this, we collected two bees from the same brood frames used to gather one-day-old bees (eight individuals in total), froze them, and subsequently analyzed their spore levels using qPCR.

### DNA extraction and sample preparation for qPCR

For DNA extraction, the abdomens of individual bees were cut using sterile instruments and placed into 2 ml cryovials, each containing 800 µl dH_2_O. The samples were homogenized using a Bead Ruptor ELITE homogenizer (Omni International) with a combination of “small” (0.5 mm) and “big” (2.8 mm) ceramid beads (Omni International).

From each homogenate, a 200 µl aliquot was taken for DNA extraction following a solution-based protocol with Nuclei Lysis Solution and Protein Precipitation Solution (Promega). The resulting DNA pellets were resuspended in 100 µl of 1× TE buffer for further analysis.

Extraction blanks (N=7) were included in each DNA extraction batch to account for possible cross-contamination at this stage of sample processing.

### Quantification of *N. ceranae* load by qPCR

To quantify the parasite load, we amplified a 65 bp fragment of the *N. ceranae Hsp70* gene using primers designed by Cilia et al. (2018). Since the *Hsp70* gene exists as a single copy per spore, the measured copy number can be directly translated into the spore load per individual (Cilia et al. 2018; Cilia et al. 2020).

The original DNA extracts were diluted 10× for qPCR which was performed using a CFX96 Touch Real-Time PCR Detection System (Bio-Rad) in a 20 µl reaction mix containing: 5 µl of DNA template, 10 µl of SsoAdvanced^TM^ Universal SYBR^®^ Green Supermix (Bio-Rad), target primers at a final concentration of 0.2 µM (see **Błąd! Nie można odnaleźć źródła odwołania.** for primer sequences), and ddH_2_O. The cycling conditions were as follows: an initial denaturation at 98 °C for 3 minutes, followed by 40 cycles of 98 °C for 15 seconds and 60 °C for 30 seconds.

For absolute quantification, a purified PCR product of a broader 824 bp *Hsp70* fragment was used as a standard. The fragment was amplified using previously designed primers (Ostap-Chec et al. 2025, see **Błąd! Nie można odnaleźć źródła odwołania.**) in a 25 µl reaction mix containing: 2 µl of DNA extract from an infected bee, 12.5 µl of DreamTaq^TM^ Hot Start PCR Master Mix, 0.4 µM of each primer, and ddH_2_O to a final volume of 25 µl. The cycling conditions were: initial denaturation at 95 °C for 3 minutes, followed by 40 cycles of 95 °C for 30 seconds, 58 °C for 30 seconds, and 72 °C for 60 seconds, with a final elongation at 72 °C for 5 minutes.

To increase yield, products from multiple reactions were pooled and subjected to agarose gel electrophoresis. The target band was then excised and purified using the ZymoClean Gel Recovery Kit. The concentration of the purified product was measured using a Qubit Broad Range Assay. The copy number was calculated based on the concentration and fragment length according to Qiagen guidelines (Qiagen 2011) for absolute quantification.

A standard curve for each plate was prepared using the purified product, with a dilution series covering a dynamic range of 8 logs (from 6.20 × 10^-1^ to 6.20 × 10^6^ copies). Standards were run in triplicate and samples in duplicate on each qPCR plate. The method demonstrated a sensitivity of 6.20 × 10^-1^ copies/µl (equivalent to 2,480 copies/bee or 3.39 log copies/bee), which was the lowest concentration in the dilution series with high reproducibility and strong linearity (R2 ≥ 0.98). PCR efficiencies for the standard curve between 90 and 110% were accepted. To confirm specificity, a melting curve analysis was performed at the end of each run, covering the temperature range of 65-95 °C in 0.5 °C increments, with a dwell time of 5 seconds per step.

### Calculation of spore load

To determine the spore load, the mean starting quantity of the template for each duplicate based on the standard curve was calculated. The results were expressed as the number of *Hsp70* copies/µl, which corresponds to the number of spores (following Cilia et al., 2018). Given that the DNA extract was diluted 10× for the reaction, the total DNA extract volume was 100 µl, and since the DNA extraction was performed on 1/4 of the original homogenate volume, the initial spore count was multiplied by 4,000. Finally, the values were log_10_-transformed to obtain the spore load as the log number of spores per bee.

### Statistics

All statistical analyses were performed in R (R Core Team 2024). Daily food consumption at each timepoint was calculated as the sum of intake from both feeders (one containing pure sucrose and the other containing sucrose with either 0.5% or 1% ethanol), divided by the number of live bees in the cage over these 24 hours. This value was expressed as per capita consumption (mg/bee). To analyze food consumption over time, we fitted a Generalized Additive Mixed Model (GAMM) using the *gamm* function from the *mgcv* package (Wood 2017; Wood 2023). The model assumed a Gaussian distribution with an identity link function and included group (control vs. infected), diet (0.5% EtOH vs. 1% EtOH), and their interaction as fixed effects. To account for potential nonlinear trends in consumption over time, a smooth term for timepoint was included, with separate smoothing functions for each group-diet combination. The smooth terms were modeled using penalized regression splines with five basis functions (k = 5). Random effects for colony and cage nested within colony were included. The results were visualized using the *ggplot2* package (Wickham 2016).

The proportion of ethanol-spiked food consumed per bee at each timepoint was calculated as the intake from the ethanol-spiked feeder per bee divided by the total food consumed per bee. This proportion ranged from 0 (no ethanol-spiked food consumed) to 1 (only ethanol-spiked food consumed). To assess changes in the proportion of ethanol-spiked food consumed over time, we fitted a Generalized Linear Mixed Model (GLMM) using the *glmmTMB* function from the *glmmTMB* package (Brooks et al. 2017). The model assumed a beta distribution with a logit link function, which is appropriate for proportion data bounded between 0 and 1. Fixed effects included timepoint (1-5), group (control vs. infected), diet (0.5% EtOH vs. 1% EtOH), and their two-way interactions. The third-way interaction was removed as it was non-significant (p = 0.294), and its removal improved model performance based on AIC comparison. Random effects for colony and cage nested within colony were included. In addition, a dispersion formula was incorporated to adjust for overdispersion. The results were visualized using *ggplot2* (Wickham 2016).

For mortality analysis, we used Cox mixed-effects regression with the *survival* package (Therneau 2022; Therneau 2023). The model included group (control vs. infected), diet (0.5% EtOH vs. 1% EtOH), and their interaction as fixed effects, with colony and cage nested within the colony as random effects. Survival curves were visualized using *ggsurvplot* from the *survminer* package (Kassambara et al. 2021).

The effect of dietary treatment on *N. ceranae* spore load in the infected group (N=80) was analyzed using linear quantile mixed model, estimating median (*lqmm* function from the *lqmm* package (Geraci 2014)), with log mean quantity of *Nosema* spores per bee as response variable, diet as fixed effect, and colony as random effect. Cage was removed from the model as it did not account for any variance and caused a singularity error. The spore load was visualized using *ggplot2* (Wickham 2016).

## Results

The analysis revealed that neither group nor diet had a significant effect on total food consumption in honeybees (group: estimate ± SE = -0.0002 ± 0.001, t = -0.179, p = 0.858; diet: estimate ± SE = -0.0008 ± 0.001, t = -0.850, p = 0.396; interaction: estimate ± SE = 0.0007 ± 0.001, t = 0.469, p = 0.639). Food consumption followed a significantly non-linear pattern across all group-diet combinations (p < 0.001 for all smooth terms; Fig. 1). The degree of non-linearity differed among combinations, with the highest non-linearity in the control group under the 0.5% ethanol diet (edf = 3.51, F = 56.92, p < 0.001) and the lowest in the control group under the 1% ethanol diet (edf = 2.78, F = 104.37, p < 0.001). In all cases, food consumption increased during the first three timepoints and then stabilized. The model explained 58.5% of the variance (R² = 0.585).

**Fig. 1.**
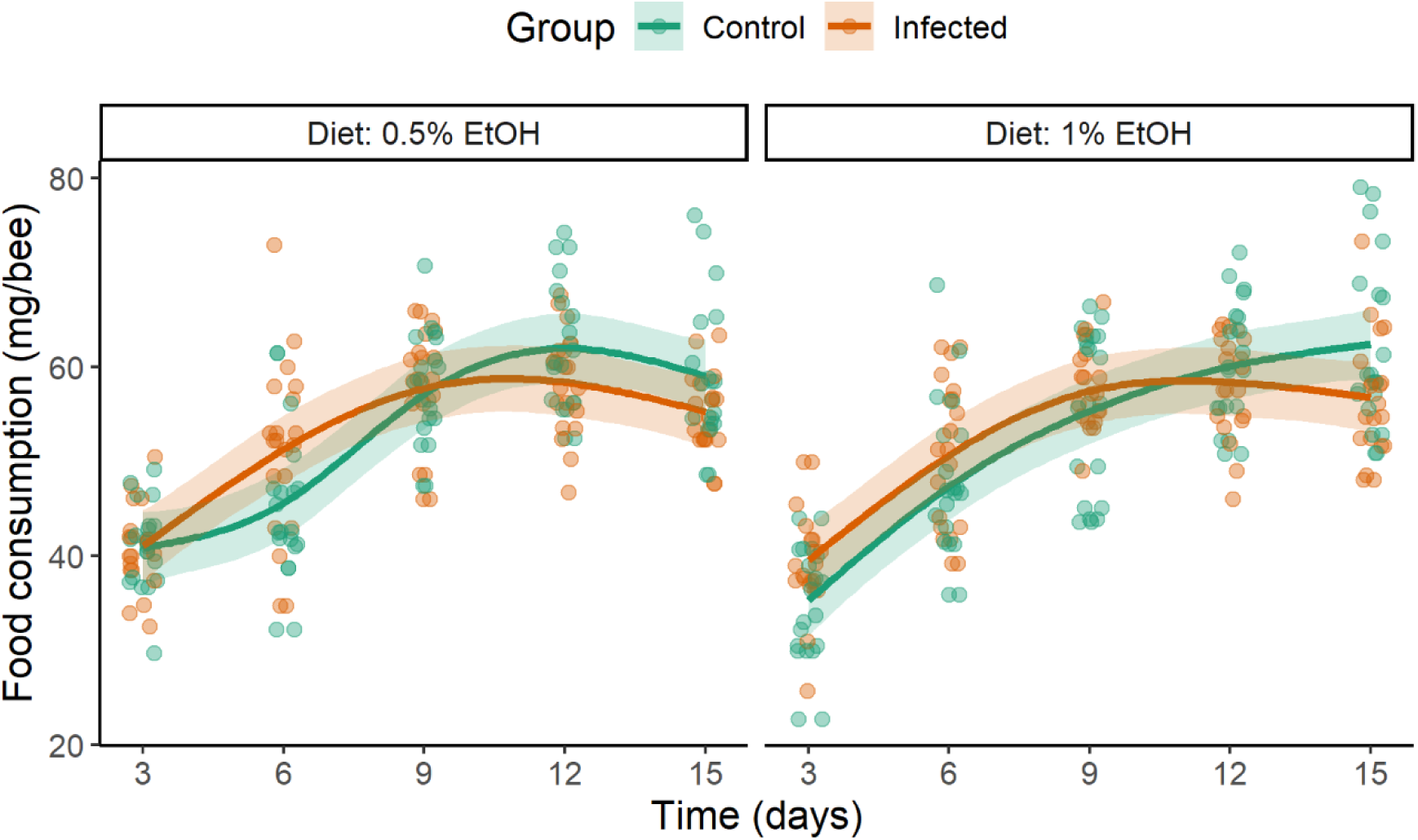
Per capita food consumption (mg/bee) in control and infected bees across five timepoints (measured every three days during 15 days) for two dietary conditions (0.5% EtOH diet vs. 1% EtOH diet). No significant differences were observed either between groups or between diets. Data points represent raw individual observations, while lines indicate GAMM predictions, and shaded areas indicate 95% confidence intervals.

Infection had a significant effect on the proportion of ethanol-spiked food consumed by honeybees (estimate ± SE = 0.234 ± 0.105, z = 2.228, p = 0.026). Infected bees had 26% (2.85% - 55.14%) higher odds of consuming ethanol-spiked food compared to the control group. Other factors and interactions were non-significant, except for an interaction between timepoint and diet (Table 2). Bees with access to 1% ethanol showed a trend to reduce the consumption of ethanol over consecutive timepoints (Fig. 2).

**Fig. 2.**
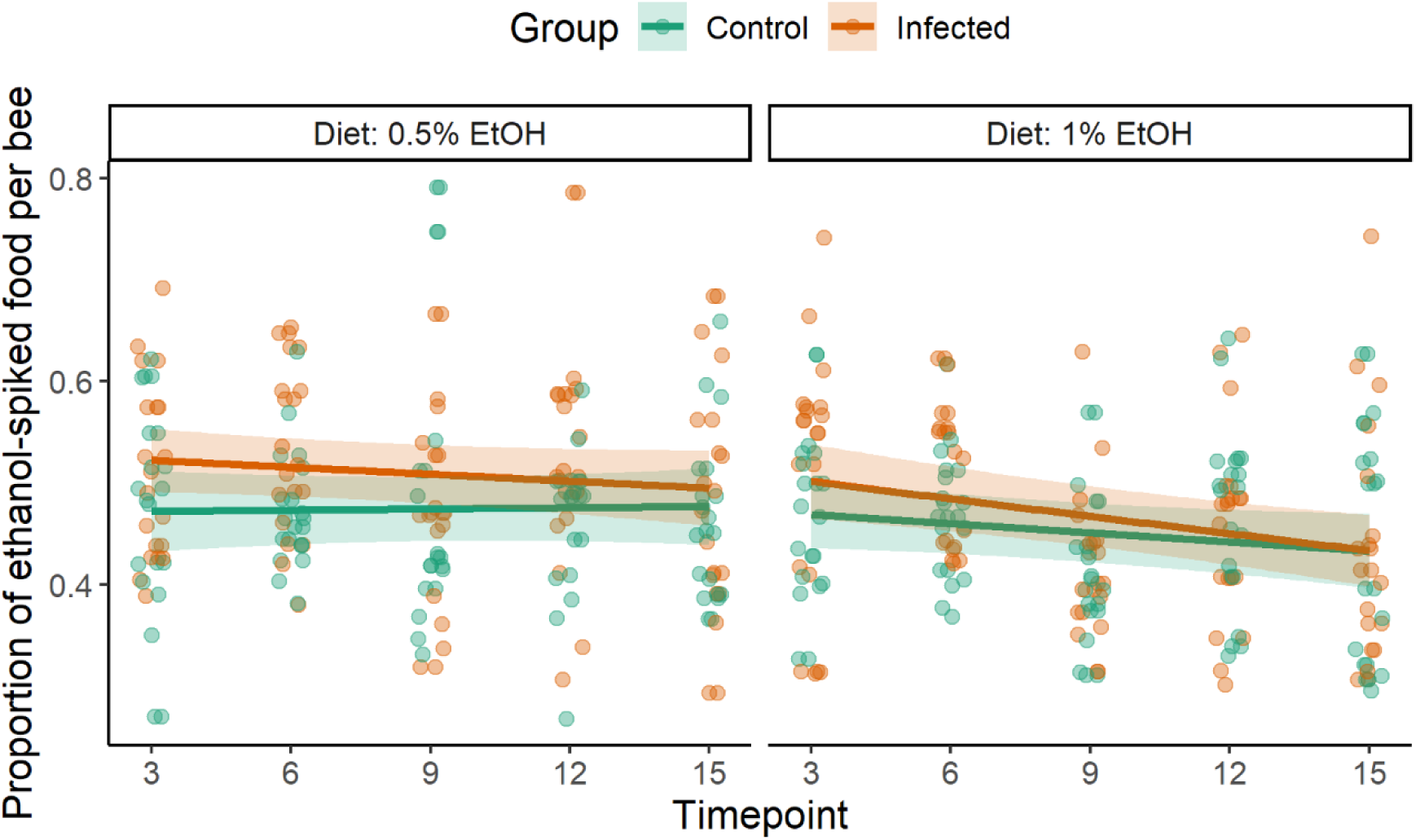
Proportion of ethanol-spiked food consumed by control and infected bees across five timepoints (measured every three days during 15 days) under two dietary treatments (0.5% EtOH diet vs. 1% EtOH diet). The proportion of consumed ethanol-spiked food was significantly higher in infected bees compared to controls. Data points represent raw individual observations, while lines represent GLMM predictions, and shaded areas indicate 95% confidence intervals.

**Table 1.**
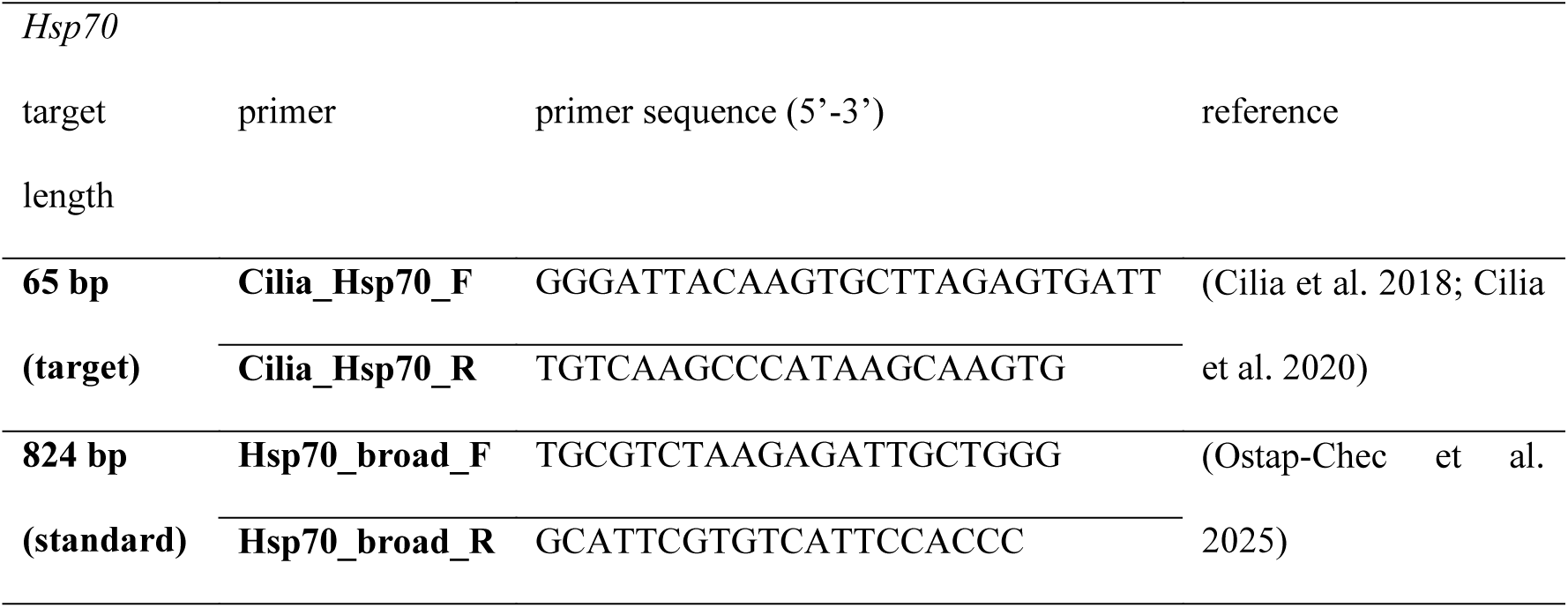
Primers used in qPCR for *Hsp70* amplification and absolute quantification.

**Table 2.** Results of the generalized linear mixed model (GLMM) analyzing the effects of timepoint, group (control vs. infected), diet (0.5% EtOH diet vs. 1% EtOH diet), and their two-way interactions on the proportion of ethanol-spiked food consumed by honeybees. The control group (uninfected bees) and the 0.5% EtOH diet were used as reference levels. Results were considered significant at p < 0.05.

| **Predictor** | **Estimate** | **Std. Error** | **z-value** | **p** |
| --- | --- | --- | --- | --- |
| **(Intercept)** | -0.11688 | 0.09733 | -1.201 | 0.2298 |
| **Timepoint** | 0.00473 | 0.02317 | 0.204 | 0.8382 |
| **Group** | 0.23366 | 0.10485 | 2.228 | 0.0259 * |
| **Diet** | 0.02917 | 0.10616 | 0.275 | 0.7835 |
| **Timepoint × Group** | -0.03216 | 0.02427 | -1.325 | 0.1851 |
| **Timepoint × Diet** | -0.04108 | 0.02413 | -1.702 | 0.0887 |
| **Group × Diet** | -0.07113 | 0.10833 | -0.657 | 0.5115 |

Mortality rates differed significantly between groups and diets. Infected bees had a 2.49-fold higher risk of death than controls (hazard ratio = 2.49, SE = 0.094, z = 9.70, p < 0.001). Bees on the 1% ethanol diet had a 26% higher risk of death compared to those on the 0.5% ethanol diet (hazard ratio = 1.26, SE = 0.104, z = 2.23, p = 0.026). However, the interaction between group and diet was significant (hazard ratio = 0.75, SE: 0.130, z = -2.17, p = 0.030), indicating that access to 1% ethanol had a weaker negative effect on survival in infected bees than in controls.

*Nosema* spore load (mean of the log mean quantity per individual ± SD) in the infected group (N=80) was 6.46 ± 1.39, over 3 orders of magnitude higher than in the control group (N=80, 3.11 ± 0.92), while the latter was comparable to background spore counts in bees sampled directly from the frames before the experiment (N=8, 3.03 ± 0.38). Spore level in extraction blanks, recalculated to the same volume as experimental samples, was 2.43 ± 1.15. The spore levels in blanks, pre-experimental bees from frames, and control group bees were negligible and consistent with previous studies (Baffoni et al. 2016; Braglia et al. 2021; Garrido et al. 2024; Ostap-Chec et al. 2025).

Comparison of *Nosema* spore load for dietary treatments in the infected group (Fig. 4) showed no significant effect of the diet (log mean quantity estimated effect ± SE = 0.3355 ± 0.2497, p = 0.19).

**Fig. 3.**
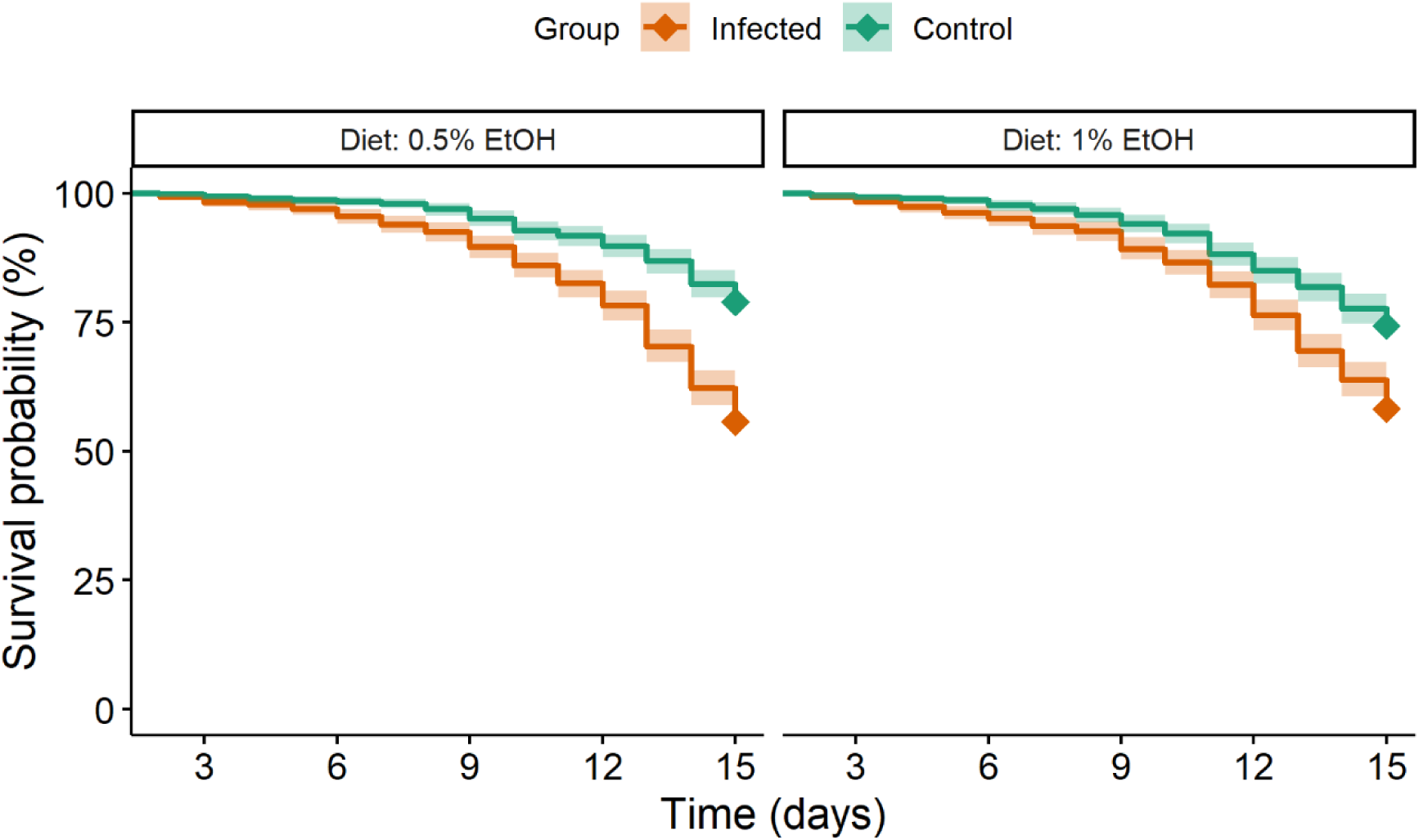
Survival plot for control and *Nosema*-infected bees over the experiment. For each replicate, 10 cages were assigned to infected bees and 10 to control bees, with five cages per diet. Each cage started with 37-40 bees. Lines represent Cox regression predictions, and shaded areas show 95% confidence intervals.

**Fig. 4.**
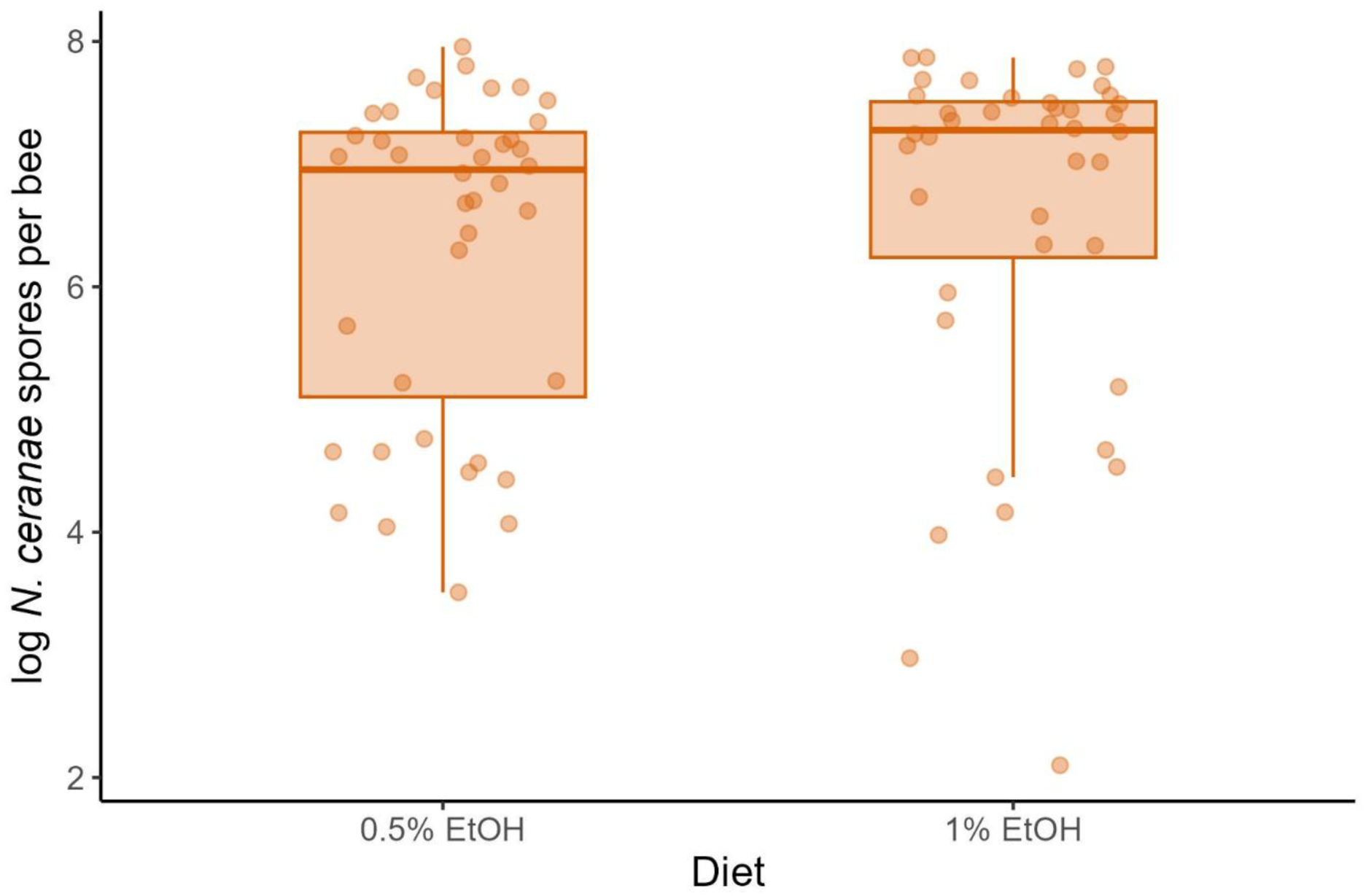
*N. ceranae* spore load per individual in the infected group for different diet treatments (0.5% EtOH vs. 1% EtOH). A single dot represents a single individual. Boxes indicate median and interquartile range, whiskers show 1.5 of the interquartile range.

## Discussion

The results of total food consumption indicate no difference between *Nosema*-infected and control bees. This aligns with our previous experimental and meta-analytic findings showing that *Nosema* infection does not affect overall carbohydrate intake (Ostap-Chec et al. 2025), supporting the idea that infected bees do not compensate for energy loss through increased sugar consumption. It is important to emphasize, however, that under natural conditions, bees have access to a more diverse diet. Energetic stress in *Nosema*-infected bees likely involves not only altered sugar metabolism but also disruptions in protein metabolism and substantial lipid depletion (Aliferis et al. 2012; Badaoui et al. 2017; Li et al. 2018; Li et al. 2019), with lipids potentially serving as an alternative energy source during infection. Nevertheless, our findings reinforce the conclusion that, at least in laboratory conditions, carbohydrate intake remains unchanged during infection, despite increased energetic demands (Mayack and Naug 2009; Mayack and Naug 2010; Holt et al. 2013).

Moreover, bees did not differ in total food consumption depending on the diet, whether it contained 0.5% or 1% ethanol. Since our ethanol-containing solutions were prepared to be iso-caloric – meaning that both pure sucrose solutions and those supplemented with ethanol had the same energy content – we can confidently state that bees on these two diets did not differ in terms of appetite. Previous studies support this conclusion. Mustard et al. (2019) found that during a 24-hour assay, bees given access to both ethanol-spiked (0.625% - 2.5%) and pure sucrose solutions consumed a similar total volume of food as control bees presented with two pure sucrose feeders. Food intake was significantly reduced only at ethanol concentrations of 5% and above. Similar patterns have been observed in other species: studies on rats and birds have shown that while high doses of ethanol can suppress appetite, low-level ethanol consumption does not affect food intake (Mazeh et al. 2008; Nelson et al. 2016). A comparable response was observed in Egyptian fruit bats (*Rousettus aegyptiacus*), which increased consumption at 0.1% ethanol, showed no effect on intake at intermediate concentrations (0.3–0.5%), but found food containing 1% ethanol or more aversive (Korine et al. 2011). These findings suggest that in species naturally exposed to ethanol in their environment, low-level, ecologically relevant concentrations are unlikely to disrupt appetite, whereas adverse effects are more likely to occur only at higher doses.

Although the overall consumption of sucrose solution did not differ between *Nosema*-infected and control honeybees, the proportion of ethanol-spiked food consumed relative to total intake revealed a pattern. Infected individuals were significantly more likely to consume ethanol-containing solutions, with 26% higher odds compared to control bees. In contrast to Mustard et al. (2019), who showed in a 24-hour assay that bees exhibit a dose-dependent preference for moderate ethanol concentrations (1.25%– 2.5%), our control bees showed no general preference for ethanol, as in their case, about half of the consumed food was ethanol-spiked. However, a preference appeared in infected bees. One possible explanation for this pattern is self-medication. This is plausible, as honeybees are known to use substances from their environment to mitigate infections (Gherman et al. 2014; Pusceddu et al. 2019). According to Abbott (2014), four criteria define self-medication: (1) active use of the substance by the host, (2) negative impact of the substance on the parasite, (3) increased host fitness when infected, and (4) cost to uninfected individuals. In our study, criterion (1) appears to be met, as infected bees increased their ethanol consumption actively. Criterion (4) is supported by previous research showing that, although bees are generally highly resilient to low doses of ethanol (Ostap-Chec et al. 2024b), its toxicity becomes apparent with age, even after occasional exposure (Ostap-Chec et al. 2024c). The cost of ethanol consumption was also demonstrated in winter foragers of different ages, where even low doses increased mortality and induced specific changes in DNA methylation (Rasmussen et al. 2021). Although criteria (2) and (3) were not directly tested in the current experiment, our results provide preliminary conclusions.

The premise that increased ethanol consumption may improve the fitness of infected individuals arises from our survival analysis. In our study, higher ethanol concentrations were associated with an increased risk of death. Specifically, bees on the 1% ethanol diet had a 26% higher mortality risk compared to those on the 0.5% ethanol diet, highlighting the potential costs of ethanol consumption. However, the interaction between infection status and diet revealed that the negative effect of 1% ethanol on survival was less pronounced in infected bees than in healthy controls. Indeed, in the absence of an interaction, infected bees given 1% ethanol would be expected to show ∼33% higher mortality than what was observed due to an additive effect of diet and infection status. This could be interpreted as an indication that ethanol consumption improves the fitness of infected individuals, thus partially fulfilling the third criterion for self-medication behavior. Whether ethanol has a detrimental effect on the parasite itself requires further research. In our study, analysis of spore load within the infected group revealed no significant differences between the two ethanol-containing diets, suggesting that a higher ethanol concentration (1%) does not have a stronger inhibitory effect on *Nosema* spores compared to the lower concentration (0.5%). However, while spore counts reflect the intensity of infection, they do not fully capture parasite fitness. A reduction in host mortality might still indicate therapeutic benefit, even without a detectable drop in spore load, but in other traits related to parasite fitness, such as transmission potential, infectivity, or spore viability. This issue requires extensive further study.

Taken together, our results suggest that infected bees are more likely to consume ethanol-containing food than uninfected individuals, and that a 1% ethanol diet may improve their survivability. While self-medication remains a plausible explanation, it is not conclusive. An alternative hypothesis is that the parasite manipulates host behavior to its benefit, a phenomenon observed across diverse host–parasite systems (Hughes et al. 2012). Parasites can alter host behavior in ways that enhance their fitness by promoting transmission, survival, or reproduction, often through effects on the nervous system or neuromodulatory pathways (Adamo 2003; Libersat et al. 2009). Striking examples include worms that cause crickets to jump into water, enabling the parasite’s aquatic life stage (Thomas et al. 2002), and the so-called “zombie ants,” which die in locations ideal for parasite development and dispersal (Hughes et al. 2011). It is therefore conceivable that *N. ceranae* may also manipulate honeybee behavior to increase ethanol consumption, potentially gaining some yet unidentified benefit. Notably, *Nosema* infections are already known to affect honeybee behavior in multiple ways. Some of these behavioral changes have been interpreted as forms of social immunity, aimed at reducing pathogen transmission within the colony. For example, infected bees often exhibit prolonged foraging bouts and reduced flight frequency and efficiency, spending more time outside the hive than uninfected individuals (Dussaubat, Maisonnasse, et al. 2013; Alaux et al. 2014; Naug 2014; Wolf et al. 2014). Similarly, the acceleration of age polyethism – where young bees begin foraging earlier than usual – has been suggested to help remove infected individuals from the social core of the colony (Lecocq et al. 2016). Although these behavioral changes may indeed function as social immunity, it cannot be excluded that they also represent a form of parasite manipulation aimed at enhancing spore transmission in the environment. In fact, there are clearer examples of such manipulation: infected bees show increased walking activity and elevated rates of trophallaxis – patterns that may actively facilitate the spread of *Nosema* spores (Lecocq et al. 2016). In addition, proteomic analyses indicate that *N. ceranae* alters host functions related to energy metabolism, immunity, and protein regulation, effectively modifying the midgut environment to enhance nutrient availability and suppress host defenses – another potential example of parasite-driven manipulation (Vidau et al. 2014). Thus, the possibility of active manipulation by the parasite cannot be excluded, and further research is needed to determine if and how ethanol affects *N. ceranae* spores, and whether increased consumption of ethanol-spiked food may ultimately benefit the parasite.

Another explanation for our findings is that ethanol primarily affects honeybee gut physiology rather than acting directly on the spores. Gut conditions strongly influence pathogen proliferation, and in honeybees, even slight disruptions to the peritrophic matrix, a key protective barrier in the midgut, can accelerate *N. ceranae* development and spore production (de Oliveira et al. 2023). Ethanol represents a complex dietary component, acting simultaneously as a toxin (Ostap-Chec et al. 2024c), a behavioral modulator (Abramson, Place, et al. 2004; Abramson et al. 2005; Bozic et al. 2006; Maze et al. 2006; Ammons and Hunt 2008; Mustard et al. 2008; Mixson et al. 2010; Wright et al. 2012; Giannoni-Guzmán et al. 2014; Black et al. 2021; Ahmed et al. 2022) and a nutritional source (Pecsenye et al. 1994; Castro et al. 2012), exerting context-dependent effects on host physiology and immunity through both direct mechanisms and modulation of the gut microbiome. In *D. melanogaster*, while low-dose ethanol ingestion decreases lifespan, it simultaneously improves intestinal barrier integrity, modulates immune pathways such as the Imd pathway, and induces metabolic shifts in the fat body, with these effects being strongly microbiota-dependent (Chandler et al. 2022). Similarly, in rodents, long-term low-dose ethanol intake has been associated with improved metabolic function, resistance to obesity, and reduced inflammation (Justice et al. 2019; Diao et al. 2020; Xuan et al. 2022). Thus, the physiological outcomes of ethanol exposure may depend on the organism’s condition, particularly under stress. Moreover, the gut microbiome, crucial for immunity, mediates the effects of various substances on immune responses. Recent studies have shown that dietary interventions, including probiotic mixtures and phytochemical supplements, can modulate the gut microbiota, enhance digestive metabolism, and strengthen immunity in honeybees (Geldert et al. 2021; Robino et al. 2024; Sbaghdi et al. 2024). Moreover, the native microbiota itself stimulates the production of antimicrobial peptides, priming bees against pathogenic infections (Kwong et al. 2017), while natural constituents such as p-coumaric acid and other plant-derived compounds have been shown to induce detoxification and immune genes and reduce pathogen loads (Boncristiani et al. 2012; Mao et al. 2013). These findings suggest that ethanol ingestion might alleviate infection-related gut dysfunction and modulate the gut environment by altering microbial composition and host metabolism, ultimately influencing the course of *Nosema* infection independently of spore load. It is also possible that the beneficial effects of low-dose ethanol are strongly dependent on the presence and stability of the gut microbiota and may only become apparent under conditions of physiological stress, such as infection.

In summary, *N. ceranae* infection does not affect overall food intake in honeybees, but infected individuals consume significantly more ethanol-spiked solutions than healthy bees. Moreover, ethanol intake may partially improve the survival of infected bees despite its general toxicity to healthy individuals, suggesting a potential self-medication effect. Our results are the first to demonstrate a prolonged preference for ethanol, potentially benefiting either the host or the parasite. However, spore load analyses showed no difference between ethanol concentrations, indicating that ethanol may not affect the parasite directly or that both doses exert similar effects. Alternatively, increased ethanol intake could result from parasite-induced manipulation or affect gut physiology and host immunity, potentially through direct or indirect modulation of the gut microbiota. Nonetheless, our findings provide new insights into honeybee interactions with bioactive compounds present in their diet and support the view that ethanol is not only a dietary component but also a behaviorally relevant and actively utilized substance. Honeybees appear capable of modulating their intake of naturally occurring compounds like ethanol in ways that may influence their physiology, behavior, and resilience to infection. Moreover, our findings suggest that ethanol plays a more significant role in honeybee ecology and behavior than previously recognized.

